# GESTURE: unsupervised genotype-specific behavioral phenotyping in rodents via graph-based representation learning

**DOI:** 10.64898/2026.08.07.743267

**Authors:** Renxiang Qiu, Sadaf Farkhani, Taha Janjua, Ece Canko, Kate Pedersen, Francois Gastambide, Ali Basirat, Christian Laut Ebbesen, Ulrike Richter

**Affiliations:** Max Planck Institute for Human Cognitive and Brain Sciences, Leipzig, Germany; Data Acquisition Systems and Analytics, Neuroscience, Lundbeck A/S, Valby, Denmark; Data Science, Global Clinical Development, Lundbeck A/S, Valby, Denmark; Symptom Biology, Neuroscience, Lundbeck A/S, Valby, Denmark; Center for Language Technology, University of Copenhagen, Copenhagen, Denmark

## Abstract

The automated quantification of complex animal behavior is fundamental to neuroscience and pharmacology, yet converting high-dimensional pose data into reproducible and interpretable behavioral measures remains challenging. Here, we present GESTURE, an unsupervised, graph-based deep generative framework that discovers and quantifies behavioral structure from pose dynamics without genotype labels or manual behavioral annotations. Applied to mice with motor dysfunction and wild-type controls, GESTURE identified a shared vocabulary of behavioral motifs. Differential use of these motifs yielded distinct “behavioral fingerprints” that reliably separated genotypes. By modeling behavior as a sequence rather than a static partition, GESTURE also quantified its temporal organization. Affected animals maintained motifs for longer and transitioned between them more predictably, indicating a slowing and stereotyping of behavioral sequences rather than simply reduced activity. GESTURE’s graph-based representation enables training across multiple recordings and embeds behavior in a shared latent space, supporting consistent cross-animal comparisons and future cross-experiment alignment. Automatically derived measures of behavioral divergence tracked the temporal dynamics of expert-annotated disability scores, reaching agreement comparable to that of independent human raters. Node- and edge-level explainability analyses further indicated that the latent representations emphasize the animal’s core motor scaffold in a biologically plausible manner. These findings support GESTURE as an interpretable, scalable framework for automated behavioral phenotyping that links genetic perturbation to quantitative behavioral phenotypes.

**Author summary:** How can behavioral changes caused by a disease-associated mutation be measured automatically and consistently? Manual assessment is slow, difficult to scale, and can vary among observers. We studied mice carrying a mutation in a calcium-channel gene whose human counterpart is associated with episodic ataxia and related movement disorders. We developed GESTURE, which learns recurring movement patterns from the tracked coordinates of a freely moving mouse without genotype labels or manual behavioral annotations. By representing the body as a set of connected landmarks, GESTURE tracks how their configuration changes over time and discovers a shared vocabulary of movements. Affected and healthy mice used this vocabulary differently. Each animal’s pattern of use formed a distinctive “behavioral fingerprint”, and these fingerprints correctly identified every animal carrying the mutation. The analysis also quantified temporal features beyond overall activity: affected mice held each movement for longer and transitioned between movements more predictably. An automatically derived severity score agreed with expert judgment as closely as two trained raters agreed with each other and yielded the same result on every run.

## Introduction

In preclinical neuroscience and pharmaceutical research, rodent behavioral assays are indispensable for assessing the therapeutic potential and safety of experimental compounds [1, 2]. Behavioral phenotyping, however, often relies on either manual scoring or coarse heuristic measures such as distance traveled or time spent mobile [3]. Manual scoring is labor-intensive, difficult to scale, and subject to inter-rater variability, whereas coarse measures discard much of the temporal and anatomical structure contained in pose data. This limitation has motivated computational approaches that combine advances in machine vision with unsupervised learning to derive automated, quantitative representations of animal behavior [3, 4].

The biological importance of this challenge is exemplified by the tottering mouse model, which carries a spontaneous mutation in the *Cacna1a* gene encoding the *α*1A subunit of the P/Q-type voltage-gated calcium channel. These channels are highly expressed in cerebellar Purkinje cells, which play a central role in motor coordination. In humans, mutations in the orthologous gene *CACNA1A* cause several neurological channelopathies, including episodic ataxia type 2 [5, 6]. Characterizing the behavioral phenotype of the tottering mouse therefore provides a translationally relevant window into genotype–phenotype relationships and may advance our understanding of human neurological disorders.

Computational ethology, enabled by advances in machine vision, has begun to address this challenge [7–9]. Early work showed that continuous behavior could be decomposed without supervision into a discrete repertoire of stereotyped units, or ‘motifs.’ MotionMapper applied wavelet transforms to video projections of freely moving fruit flies to generate a 2D map of behavior [10], whereas MoSeq used principal component analysis (PCA) of depth-camera data to segment mouse behavior into ‘syllables’ organized in a grammatical structure [11]. Two features of these early approaches limit their broader application. First, several influential pipelines were tied to specific sensing modalities, particularly depth-camera representations [11, 12], restricting their transfer to widely used video-based pose-estimation workflows. Second, many relied on linear dimensionality reduction. This approach is well suited to a simple body plan, for which four principal components account for most postural variance in *C. elegans* [13]. However, it is less naturally suited to the coordinated, nonlinear dynamics of animals with articulated limbs.

Deep learning-based tools for markerless pose estimation, including DeepLabCut [4, 14] and SLEAP [15], now enable the extraction of precise 2D or 3D coordinates for user-defined anatomical keypoints from standard video. These coordinates provide a high-resolution view of posture over time and have shifted behavioral analysis from low-dimensional summary measures toward high-dimensional pose time series. This development has also created a new analytical challenge: how to distill meaningful and interpretable behavioral states from raw coordinates that are high dimensional, temporally dependent, and variable across animals and sessions [3].

A range of unsupervised methods has since been developed for these pose outputs. One family, exemplified by B-SOiD [16], first constructs kinematic features such as joint velocities and angles and then clusters them in a nonlinear embedding. Although interpretable, this approach relies on a priori assumptions about which features are behaviorally relevant and may overlook more holistic patterns. A second family, including VAME [3], BehaveNet [17], and keypoint-MoSeq [18], learns directly from pose trajectories and captures temporal structure more flexibly. However, these methods represent the body’s spatial organization only implicitly through concatenated coordinates or latent pose states. More recently, graph neural networks have been used to encode body parts as nodes and their couplings as edges [19–22]. This formulation is well suited to articulated pose, in which behavioral meaning depends on coordinated relationships among body parts rather than isolated point trajectories, and provides a more direct link between learned features and anatomy. Three challenges remain: capturing pose-dependent relational changes between body parts in an interpretable manner, even when a fixed anatomical graph provides the scaffold; explaining what these models learn to support biological interpretation; and avoiding dataset-specific representations that prevent behavior from different animals or experimental conditions from being embedded in a shared latent space for direct comparison.

To address these challenges, we introduce **GESTURE** (**G**raph-based **E**dgeConv **S**patio-**T**emporal **U**nsupervised **R**odent **E**mbedding), a framework designed to balance expressive nonlinear modeling with biological interpretability. GESTURE combines three components: (1) window-wise egocentric alignment and a fixed anatomical graph whose edge features are computed dynamically from the instantaneous pose; (2) an EdgeConv-based *β*-variational autoencoder that learns anatomically structured latent spaces; and (3) a Gaussian hidden Markov model that discretizes latent trajectories into coherent behavioral motifs. The framework thus combines graph-based inductive structure with the flexibility of deep generative modeling, enabling motif discovery from learned anatomical relationships rather than exclusively from preselected kinematic summaries.

We apply GESTURE to the unsupervised phenotyping of tottering and wild-type mice. The framework discovers a shared repertoire of behavioral motifs and reveals genotype-specific ‘behavioral fingerprints’ that distinguish animals without labels. Behavioral divergence measures derived from these fingerprints also track expert-annotated disability scores, linking the model outputs to an external phenotypic measure. These results support GESTURE as an interpretable pipeline for automated behavioral phenotyping in both basic and translational neuroscience.

## Materials and methods

### Experimental model and data acquisition

All behavioral data were collected from the tottering mouse model (B6.D2-*Cacna1a*^tg^/J; JAX stock #000544) [23], a well-established model of episodic motor attacks and ataxia-like phenotypes [5, 6]. Tottering mice carry a point mutation in the *Cacna1a* gene that disrupts Cav2.1 calcium channel function, resulting in Purkinje cell dysfunction and episodic motor instability characteristic of the ataxia-like phenotype.

The study included an initial cohort of 16 age-matched male mice (8 wild-type (WT) and 8 tottering (TG)). Each animal was recorded individually for 2 h in a standardized open-field arena at 25 frames per second (1024 × 768 pixels). All experimental procedures were approved by the Danish Animal Experiments Inspectorate (Dyreforsøgstilsynet) under licence no. 2024-15-0201-01687 and conducted in accordance with Danish legislation implementing EU Directive 2010/63/EU on the protection of animals used for scientific purposes.

Following acquisition, all recordings underwent quality control. One recording was excluded because of severe overexposure, resulting in a final dataset of 15 videos (7 WT and 8 TG). Because GESTURE is trained without labels, a train–test split is not required, and the encoder could in principle be fitted to the entire corpus. Instead, we fitted it to a balanced subset of six recordings (three WT and three TG) that were free of technical artifacts such as overexposure or camera freezing, thereby preventing corrupted frames from shaping the learned representation. The subset was selected solely on the basis of recording quality; no phenotype-related information was used during fitting. The resulting encoder was then applied unchanged as a fixed feature extractor to all 15 recordings; the nine recordings not used for fitting are analyzed separately in Zero-shot generalization to unseen animals.

### Pose estimation and preprocessing

For pose estimation, we used DeepLabCut [4], which was fine-tuned for our experimental setup to track 11 keypoints (nose, head center, ears, neck, body center, four paws, and tail base) with high fidelity. To emphasize intrinsic postural dynamics rather than the animal’s absolute position or heading in the arena, we applied a window-wise egocentric alignment pipeline (Fig 1a). Here, an *analysis window* is a contiguous sequence of frames constituting a single model input sample.

**Fig 1.**
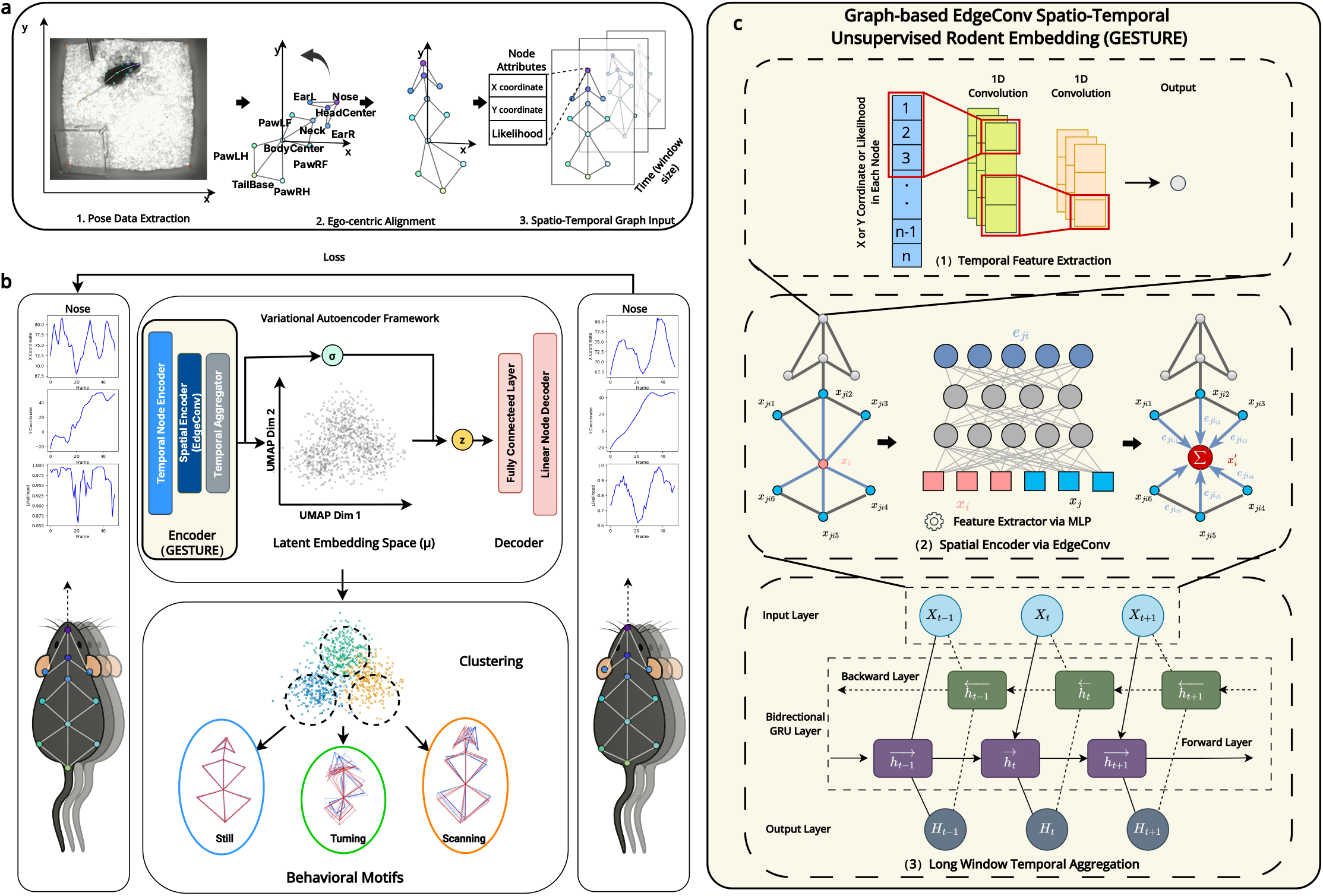
Preprocessing pipeline and GESTURE architecture. **a) Preprocessing.** Raw 2D keypoint trajectories extracted from video are converted into a standardized input format. The pipeline comprises (i) window-wise egocentric alignment through translation and rotation and (ii) representation of each pose as a graph, with keypoints as nodes and anatomical connections as edges. Keypoint labels correspond to anatomical landmarks (e.g., *Body Center* and *Neck* define the alignment reference direction). Sliding windows of these pose graphs form the final input tensor. **(b) Model architecture.** GESTURE is a *β*-variational autoencoder designed to learn a structured latent representation of behavior. The encoder uses temporal convolutions and an EdgeConv-based spatial encoder to process the input graphs. The resulting features are mapped to a latent space by predicting the parameters (*µ*, *σ*) of a Gaussian distribution, from which a latent vector (*z*) is sampled. The decoder then reconstructs the original pose sequence from this vector. The model is trained end-to-end by minimizing a combined reconstruction and KL-divergence loss. **(c) Encoder modules used in GESTURE.** (1) Temporal feature extraction: per-keypoint 1D temporal convolutions produce short-term motion features. (2) Spatial encoding (EdgeConv): edge features are computed between anatomically connected keypoints and aggregated to update node embeddings. (3) Temporal aggregation: features are aggregated by a bidirectional GRU.

For each analysis window, unreliable keypoint detections (likelihood ≤ 0.8) were removed and linearly interpolated, after which a Savitzky–Golay filter was applied to reduce high-frequency noise [24]. We first translated all keypoint coordinates so that the ‘BodyCenter’ keypoint was located at the origin. We then aligned the animal’s orientation by rotating all keypoints such that the reference direction was consistent across samples. This reference direction was defined as the *window-mean* vector from ‘BodyCenter’ to ‘Neck.’ The window-mean reference vector avoids frame-wise discontinuities while preserving temporal smoothness and ensures that similar postures are represented consistently regardless of their absolute location or orientation in the arena.

Finally, the continuous pose data were segmented into overlapping temporal windows to serve as input for the GESTURE model.

### The GESTURE framework

We developed GESTURE, a deep generative framework for unsupervised behavioral phenotyping (Fig 1b). It comprises a spatiotemporal encoder that maps high-dimensional pose sequences to a low-dimensional latent space and a decoder that reconstructs the original sequence. The encoder captures both the spatial configuration of the body and its temporal evolution.

**1. Temporal feature extraction.** For each of the 11 keypoints, a dedicated 1D temporal convolutional network extracts low-level, short-term motion features (e.g., velocity- and acceleration-like cues) independently from the coordinate and likelihood traces (Fig 1c1). This stage captures fine-grained temporal patterns within each node before any spatial interaction occurs, allowing the model to disentangle local motion dynamics from interlimb coordination. The resulting per-node temporal embeddings are then passed to the spatial encoder to learn geometric relationships across keypoints.
**2. Spatial encoding via EdgeConv.** GESTURE models spatial relationships between connected body parts using **EdgeConv** [20] (Fig 1c2). We represent the animal as an anatomical graph whose connectivity reflects anatomical adjacency, whereas the learned edge features vary with the instantaneous geometry of the pose. This approach allows the model to capture context-sensitive representations of posture and coordination without requiring handcrafted pairwise features.

Specifically, EdgeConv defines edge features that capture the relationship between a central node (keypoint) and its neighbors (Fig 1c3). For each node *i*, we define its neighborhood N (*i*) as the set of anatomically connected keypoints. For each neighbor *j* ∈ N (*i*), an edge feature **e***_ij_* is computed using an asymmetric function implemented as a multilayer perceptron (MLP):

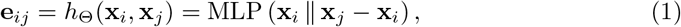

where **x***_i_* and **x***_j_* are the features of the respective nodes, and ∥ denotes feature concatenation. This formulation simultaneously captures the global position of the central node (**x***_i_*) and the relative local position of its neighbor (**x***_j_* − **x***_i_*), allowing the model to learn fine-grained geometric cues such as relative distances, directions, and angular orientations.

The computed edge features are then aggregated to update the representation of the central node. While the original EdgeConv paper proposed max-pooling, we adopt a **summation** strategy:

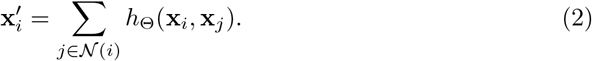

This aggregation choice allows all neighboring nodes to contribute to the final representation, preserving relational information distributed across the neighborhood. Compared with max-pooling, summation yields smoother gradients during training and can improve convergence.

**3. Temporal aggregation.** To model the temporal structure of behavior, the GESTURE encoder incorporates a recurrent aggregation module that integrates spatial features over time. Specifically, the sequence of node-level representations produced by the EdgeConv layers is processed by a bidirectional gated recurrent unit (Bi-GRU) [25, 26]. This design enables the model to capture long-range temporal dependencies across extended behavioral sequences while remaining sensitive to coordinated multijoint dynamics across consecutive frames. By explicitly modeling temporal evolution within a unified recurrent framework, GESTURE learns a temporally coherent representation of pose dynamics that supports downstream behavioral segmentation and motif discovery.

### Latent space and decoder

The encoder maps the final spatiotemporal features into a compressed latent space using a *β*-variational autoencoder (*β*-VAE) framework [27]. It outputs the mean *µ* and variance *σ*^2^ of a Gaussian distribution, from which a latent vector *z* is sampled. The model is trained to minimize the combined loss function

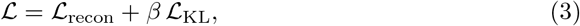

where L_recon_ is the mean squared error between the original and reconstructed pose sequences and ensures reconstruction fidelity. L_KL_ is the Kullback–Leibler (KL) divergence, a regularization term that encourages a well-structured, disentangled latent space. The hyperparameter *β* (set to 10) balances these two objectives [28]. A simple feed-forward decoder then reconstructs the pose sequence from the latent vector *z*.

### Unsupervised discovery of behavioral motifs

To discretize the continuous latent space into meaningful behavioral states, we use a Gaussian hidden Markov model (G-HMM) [29]. We model the sequence of latent vectors {*z_t_*} as being generated by a system with *K* hidden states (behavioral motifs). Each state *k* is associated with a Gaussian emission distribution N (*µ_k_,* Σ*_k_*) in the latent space. The G-HMM jointly learns the emission distributions and transition probabilities, thereby grouping similar behaviors while maintaining temporal coherence.

Windows containing corrupted input (camera freezes) are nearly identical to one another and therefore form a highly self-transitioning state that would inflate dwell-time and sequence statistics. We excluded this state from all occupancy, bout-duration, and transition analyses and renormalized each animal’s occupancy over the remaining motifs. Because deleting windows and splicing the remainder would merge motif runs that were never adjacent, we instead *cut* each recording at the gaps left by the excluded windows and treated every run as an independent sequence.

The rationale for preferring a sequence model over a static partition such as *k*-means extends beyond smoother labels. Behavioral states commonly overlap in latent space, and a method that clusters each window independently of time must resolve this overlap from position alone. To illustrate this point, we simulated data from a G-HMM with known parameters, for which the true state sequence was available as ground truth, and fitted both models to the same windows (Fig 2). Both recovered most individual windows correctly, but only the G-HMM recovered the *sequence*. It returned 19 behavioral segments when 17 were present, whereas *k*-means fragmented the same data into 122 segments. Nevertheless, *k*-means assigned 88% of individual windows to the correct state. Per-window accuracy is therefore insensitive to the relevant failure in behavioral analysis because bout durations, transition structure, and motif sequencing are properties of the segmentation rather than of individual windows.

**Fig 2.**
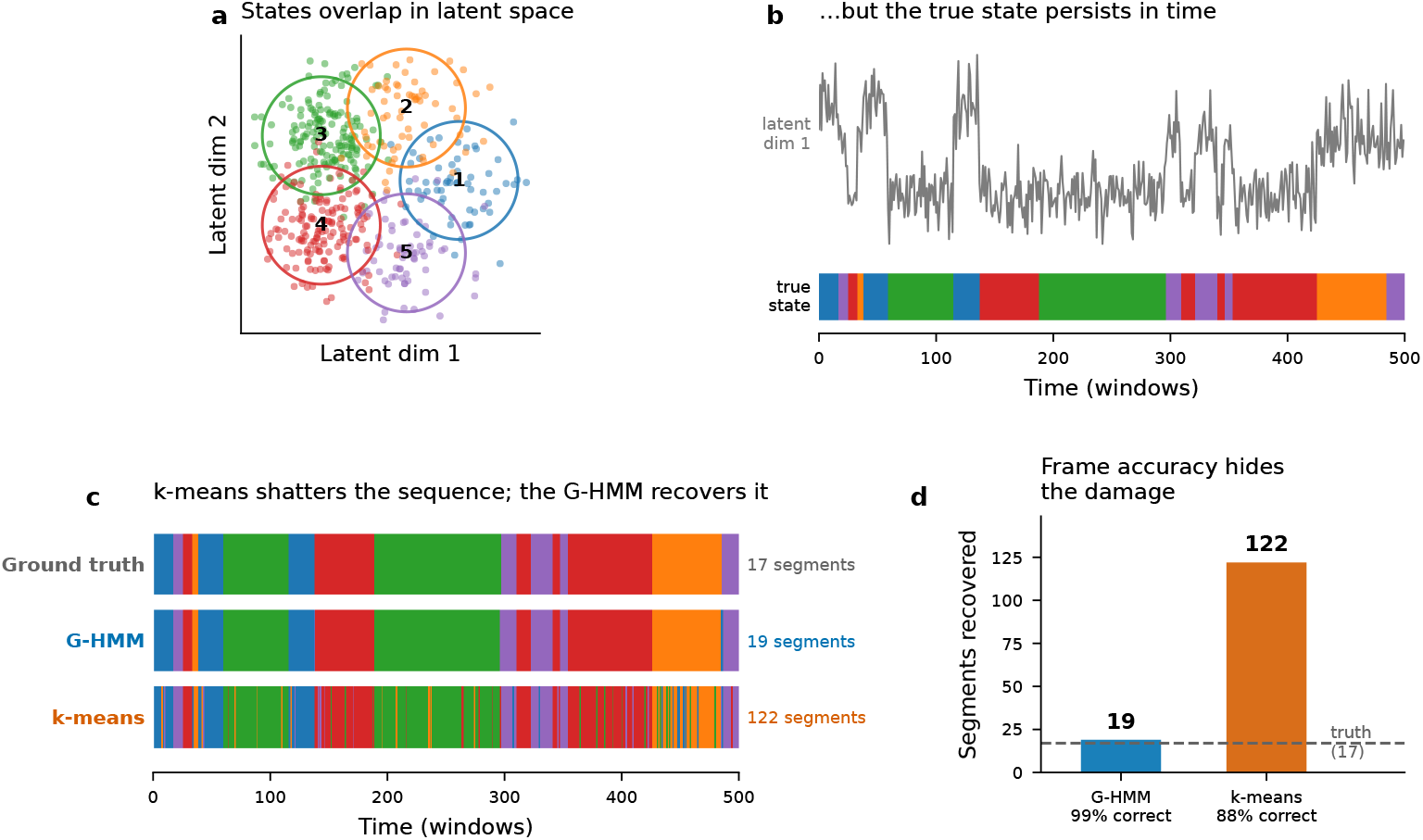
Why a Gaussian HMM rather than *k*-means: synthetic data with known ground truth. Data were simulated from a G-HMM with five states and deliberately overlapping Gaussian emissions, and a G-HMM and *k*-means were then fitted to the identical windows. **(a)** The latent space. The five emission distributions overlap substantially, so position alone is often ambiguous. Circles mark ±2*σ*. **(b)** Temporal ordering of the same data. Despite the overlap, the true state sequence persists, reflecting temporal coherence in behavior. **(c)** Recovered state sequences. The G-HMM tracks the ground truth, whereas *k*-means switches between states when their emissions overlap because it does not use preceding states. **(d)** Effect of ignoring temporal information. Although *k*-means labels 88% of individual windows correctly, it oversegments the sequence more than sevenfold (122 segments when 17 are present). Per-window accuracy does not capture this fragmentation. The transition prior of the G-HMM resolves the ambiguity and recovers 19 segments.

### Model explainability

To identify the features learned by the model, we use two explainability techniques.

#### Node importance

We use GNNExplainer [30] to identify the keypoints (nodes) that most strongly influence the model’s latent representation. GNNExplainer learns a soft mask over the nodes to identify the minimal subset that preserves the latent embedding, with higher mask values indicating greater importance.

#### Edge importance

We developed a gradient-based saliency method tailored to the geometry-aware nature of EdgeConv. We compute the gradient of the latent embedding norm (*S* = ∥ **z**∥ _2_) with respect to the node features. The magnitude of the gradient difference across an edge (*i, j*) reflects the model’s sensitivity to the relative positions of the two keypoints:

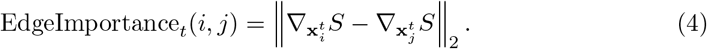

This provides a fine-grained view of which anatomical relationships are most critical for defining a given behavior.

### Model architecture and hyperparameter choices

The GESTURE model processes 11 anatomical keypoints, each represented by three features (*x*, *y*, and likelihood). The encoder combines per-node temporal convolutions, graph-based spatial reasoning through EdgeConv on an anatomical graph, and an optional bidirectional GRU block for long-range aggregation. The latent space is 32-dimensional, with the mean and variance predicted by linear layers within the variational framework. The decoder mirrors this structure and reconstructs the original spatiotemporal pose data from the latent code.

Using a window size of 50 frames, the model was trained for 150 epochs with a batch size of 32 and an initial learning rate of 8 × 10^−4^. A ReduceLROnPlateau scheduler reduced the learning rate when the validation loss plateaued (patience, 5; factor, 0.5).

## Results

### GESTURE improves reconstruction fidelity over baselines

We first evaluated the technical validity of GESTURE by assessing how faithfully it reconstructed high-dimensional pose sequences. Using the same training protocol for all models, we benchmarked GESTURE against representative baselines, including VAME [3], DeepOF [21], and a standard spatiotemporal GCN [19]. GESTURE consistently achieved lower reconstruction error and faster convergence than all baselines (Fig 3a).

**Fig 3.**
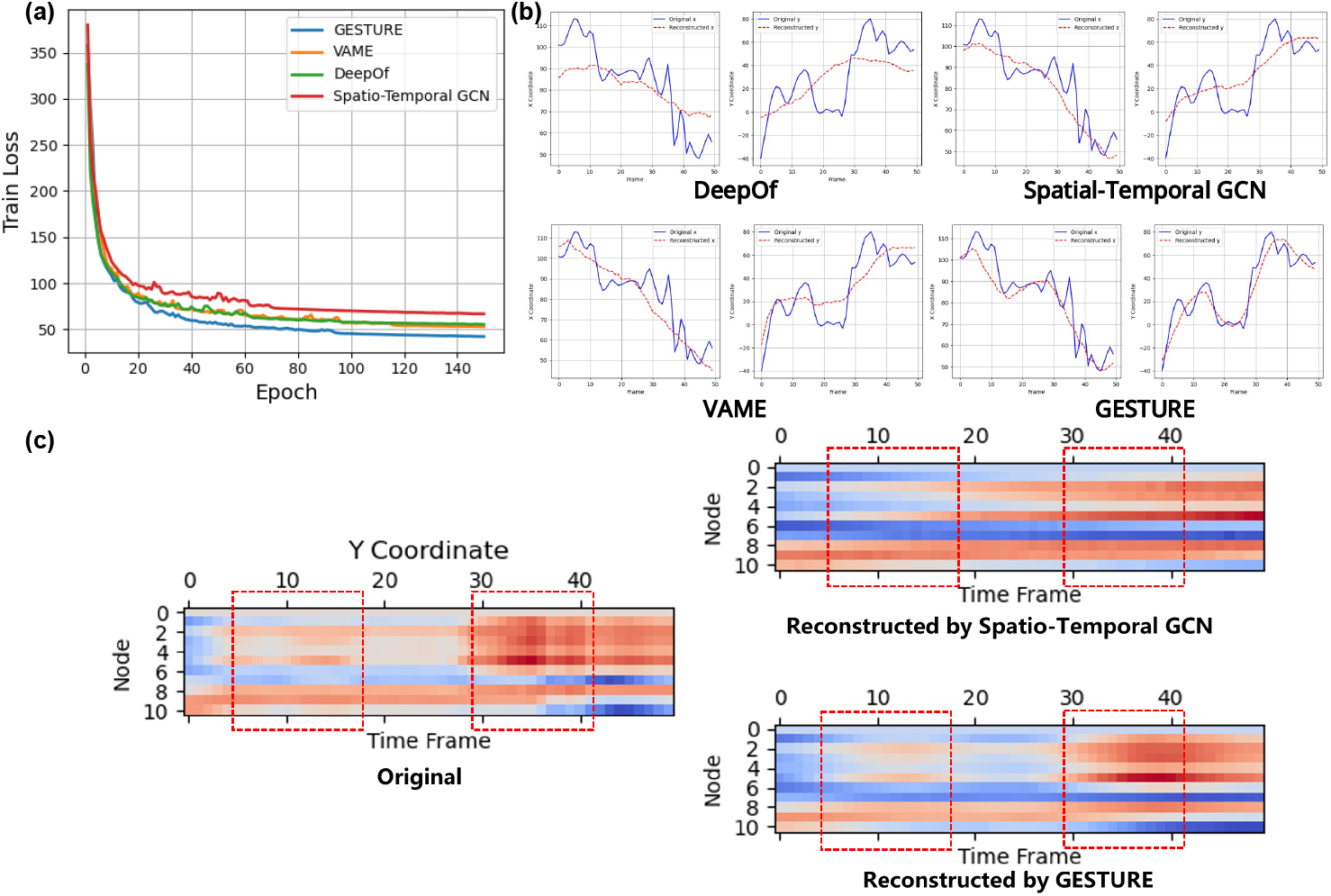
Reconstruction performance across models. **(a)** Training loss (MSE) across epochs for each model. **(b)** One-dimensional reconstruction of a representative keypoint. Blue, ground-truth trajectory; dashed red, reconstructed trajectory. **(c)** Reconstruction heatmaps of the *y* coordinates of all 11 keypoints over 50 frames. Red boxes highlight segments with pronounced dynamics captured by GESTURE but missed by the baseline models.

To assess reconstruction quality beyond the aggregate loss, we visualized representative trajectories and full keypoint heatmaps. GESTURE followed complex, nonmonotonic coordinate trajectories more accurately than the baseline models (Fig 3b) and reproduced more of the temporal structure across all 11 keypoints in the node-by-time heatmaps (Fig 3c). These results show that GESTURE more faithfully reconstructs spatiotemporal pose dynamics, a prerequisite for extracting biologically meaningful behavioral structure from video-derived postural measurements [31].

### Interpretable behavioral vocabulary

We hypothesized that the structured latent space would contain distinct clusters corresponding to a vocabulary of discrete behavioral motifs. Applying a G-HMM to the pooled latent embeddings from all 15 mice (7 WT and 8 TG) identified an optimal set of *K* = 10 states according to the BIC. These states were not animal-specific but instead constituted a shared repertoire. The average motion profile of each state supported a descriptive, ethologically meaningful label (Fig 4a). One state did not represent a behavior: cluster 0 captured windows containing corrupted input (see below) and was excluded from all subsequent comparisons. Cluster 7 was a genuine low-motion state in which the pose estimator frequently lost the tail base. Its mean within-window displacement of 0.6 px was the second lowest of the ten states.

**Fig 4.**
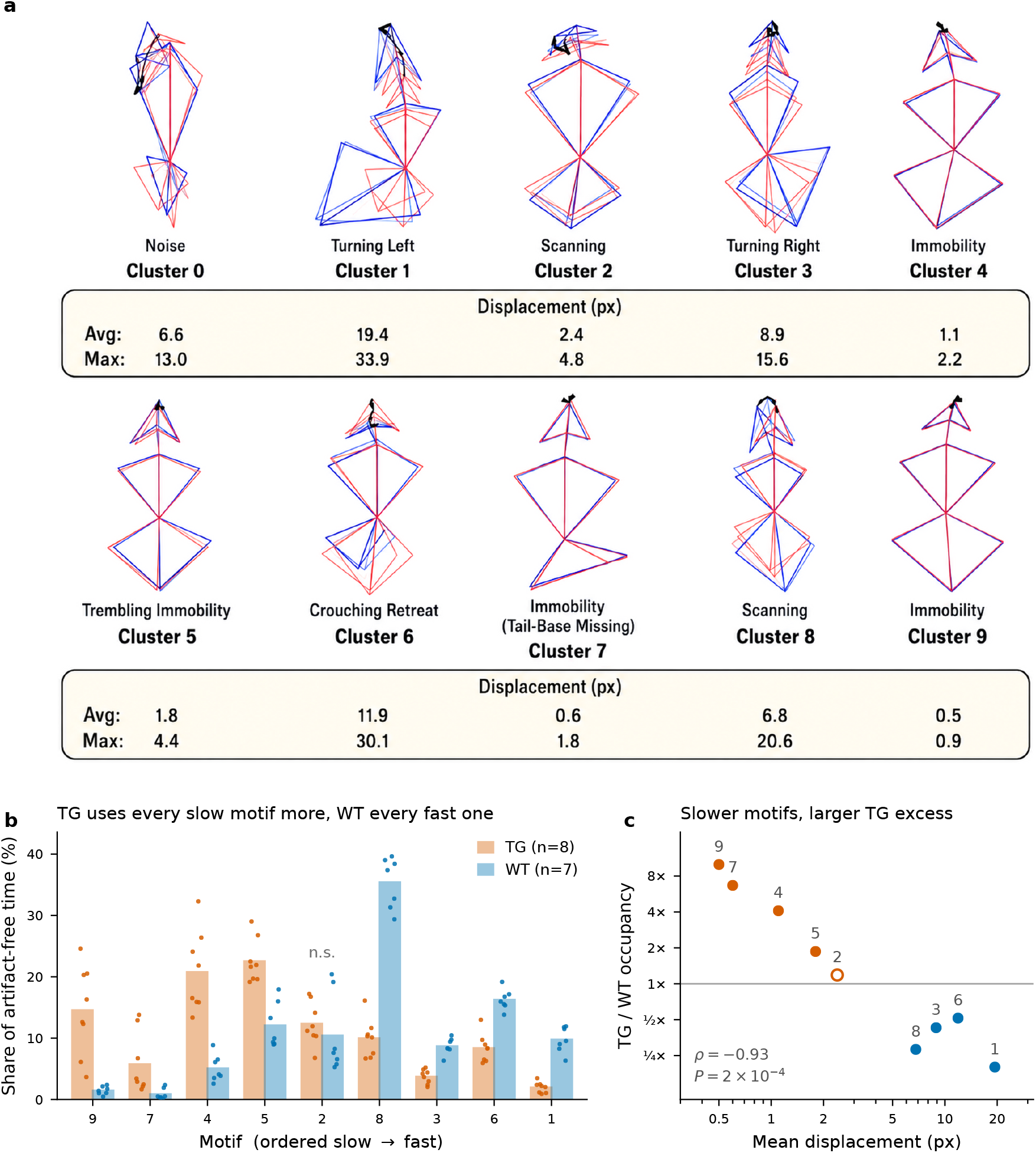
Genotypes share a behavioral vocabulary; tottering mice shift usage toward slower motifs. **(a)** Ethogram of the ten GESTURE-defined states. Panels show representative skeleton trajectories (color-coded from red to blue over 2 s), ethological labels, and mean and maximum keypoint displacements. *Note:* Trajectories are recentered on the mean body center rather than the first frame; turning movements therefore form fan-shaped arcs in which head paths may initially deviate in the direction opposite to the net turn. Cluster 0 (*Noise*) represents recording artifacts; cluster 7 is a low-motion state with frequent loss of tail-base tracking. **(b)** Proportion of session time spent in each motif, stratified by genotype. Cluster 0 is excluded, and the remaining nine motifs are renormalized to prevent bias from unequal artifact loads. Motifs are ordered from slowest to fastest by mean displacement. Bars show group means; dots represent individual animals (*n* = 8 TG, *n* = 7 WT). All motifs except one (marked n.s.) differ significantly between genotypes (Mann–Whitney test, Benjamini–Hochberg *q <* 0.05). **(c)** Ratio of mean TG occupancy to mean WT occupancy for the nine motifs, plotted against mean displacement. Colors indicate the predominant genotype; the open point denotes the nonsignificant motif in (b). Labels show the cluster indices from (a); *ρ* denotes Spearman’s rank correlation.

The two genotypes drew on the *same* vocabulary but used it differently, with the difference organized almost entirely by speed. TG animals used each of the five slowest motifs more frequently than WT animals, whereas WT animals used each of the four fastest motifs more frequently (Fig 4b; eight of the nine motifs differed at *q <* 0.05).

The shift was graded rather than categorical: across the nine motifs, the TG-to-WT occupancy ratio scaled inversely with mean within-window displacement (Spearman *ρ* = −0.93, *P* = 2 × 10^−4^; Fig 4c), ranging from a 10-fold TG excess in the slowest motif (*Immobility*, cluster 9) to a 5-fold WT excess in the fastest motif (*Turning Left*, cluster 1). Thus, the repertoire was shared, but occupancy in TG animals was systematically shifted toward the slower end of the speed axis.

### Genotype-specific behavioral fingerprints

Although TG and WT mice shared the same underlying behavioral vocabulary, their temporal deployment of these motifs revealed a clear genotype-dependent divergence. When the trained GESTURE model was applied to the full 15-video corpus (7 WT and 8 TG), the resulting Gantt charts showed that WT mice typically exhibited diverse, rapidly changing motif sequences. In contrast, TG mice spent long, uninterrupted periods in the low-activity states: the three *Immobility* variants (clusters 4, 7, and 9) and *Trembling Immobility* (cluster 5) (Fig 5a). This difference is quantified below.

**Fig 5.**
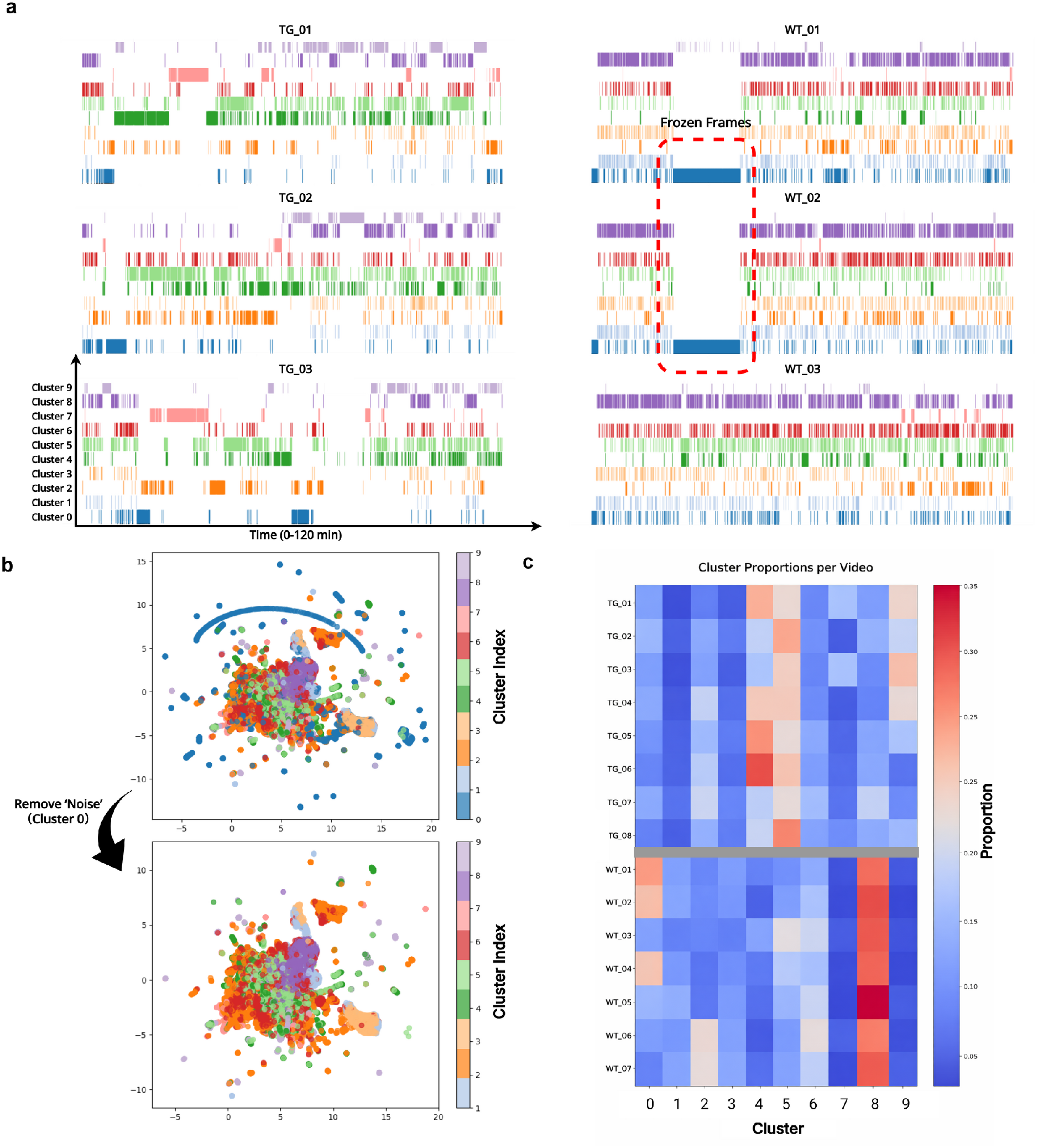
GESTURE generalizes to the full dataset and separates behavioral structure from recording artifacts. **(a)** Representative Gantt charts for six videos (3 TG, left; 3 WT, right). WT mice exhibit diverse, rapidly changing behavioral sequences, whereas TG mice display long, uninterrupted blocks of low-activity states. Some WT sessions contain camera-freeze artifacts, which appear as solid segments of *cluster 0*. **(b)** UMAP projection of all 2-s windows. Top, windows colored by cluster label, with a linear manifold corresponding to frozen-frame outliers; anomalies are isolated in *cluster 0*. Bottom, embedding after removal of *cluster 0*. **(c)** Behavioral fingerprints (cluster-occupancy distributions) for all 15 valid videos.

Extending the analysis to the full dataset also revealed recording-related artifacts. Three WT sessions contained frozen-frame segments caused by camera lockups. In a uniform manifold approximation and projection (UMAP) embedding [32] of the latent windows, these segments formed a distinct linear manifold and were assigned to a dedicated *cluster 0*, separate from the manifold occupied by genuine behavioral motifs (Fig 5b).

Behavioral fingerprints computed across the 15 valid videos showed the same cohort-level divergence: WT animals distributed their occupancy across a broader set of motifs, whereas TG animals concentrated their occupancy in a smaller set of low-activity motifs (Fig 5c).

To quantify this pattern, we defined each animal’s behavioral fingerprint as the probability distribution of time spent in each of the 10 discovered states. TG mice spent a larger fraction of time in the *Immobility* motif, whereas WT mice distributed their occupancy more evenly across active and inactive states (Fig 6a, left). Pairwise **Hellinger distances** between fingerprints produced a pronounced block-diagonal matrix with low within-genotype distances and high between-genotype distances (Fig 6b).

**Fig 6.**
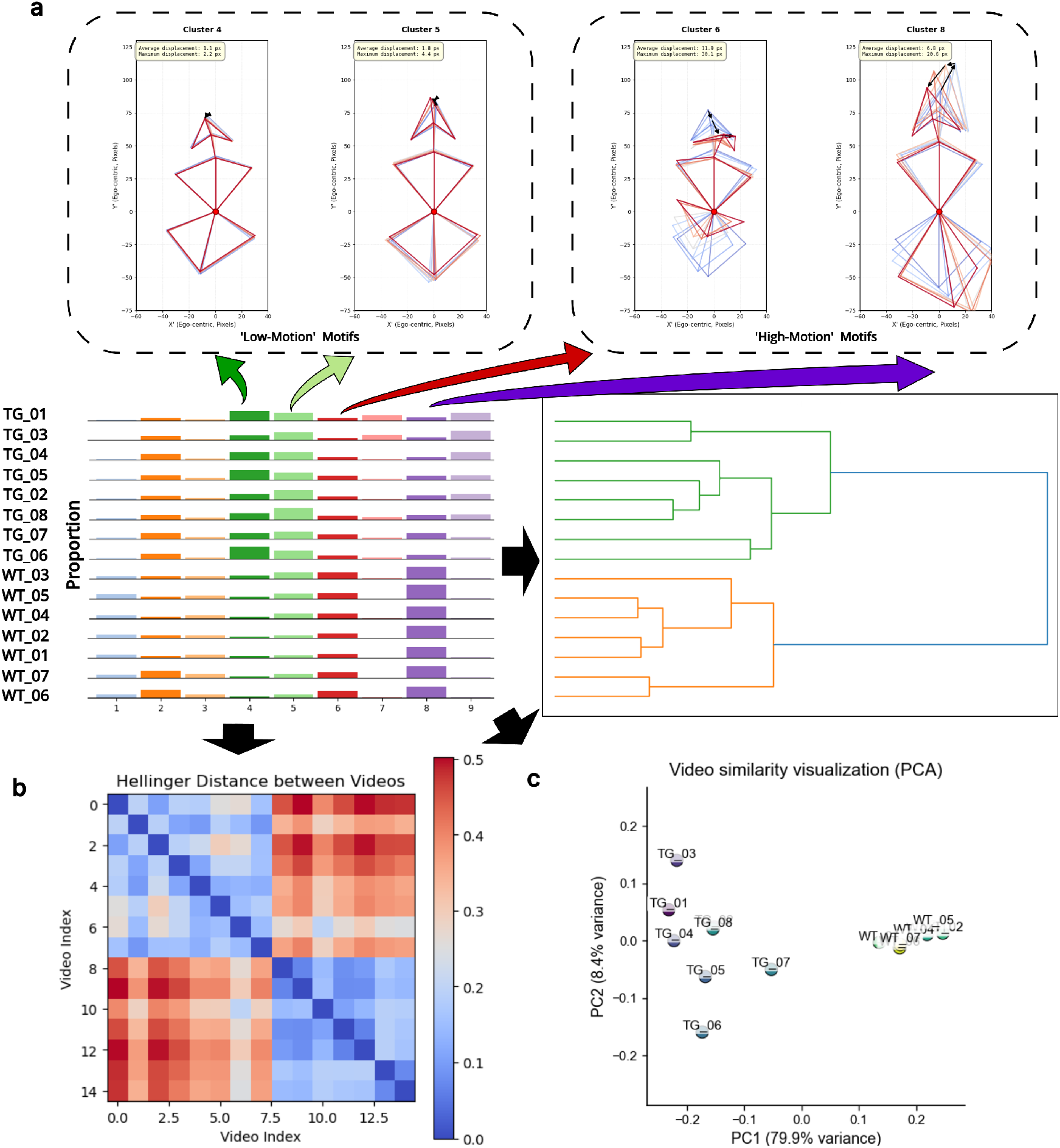
Behavioral fingerprints enable unsupervised separation of genotypes. **(a)** Top, representative motifs for clusters 4, 5, 6, and 8. Bottom left, behavioral fingerprints (motif occupancy) for each mouse. Bottom right, a dendrogram obtained by hierarchical clustering of these fingerprints separates WT from TG mice. **(b)** Hellinger-distance heatmap showing low within-genotype distances (blue) and high between-genotype distances (red). **(c)** Principal-component projection of the fingerprints. PC1 (79.9% of the variance) separates the genotypes without overlap, and its loadings recover the motif speed axis; PC2 (8.4%) is unrelated to speed and captures heterogeneity within the TG group. Loadings are reported in the text. Statistical validation of this separation is shown in Fig S3 Fig.

Using these distances, we performed agglomerative **hierarchical clustering** (average linkage) to group animals solely by behavioral similarity. This entirely unsupervised procedure produced a dendrogram in which the two primary branches corresponded exactly to the WT and TG groups, with no misassignments (Fig 6a, bottom right). A leave-one-out nearest-neighbor classifier likewise recovered genotype from the fingerprint alone in all 15 animals, and the separation remained unchanged after removal of the artifact state (Fig S3 Fig).

Principal-component analysis of the same fingerprints made the structure of this separation explicit (Fig 6c). One component dominated: PC1 accounted for 79.9% of the between-animal variance and alone separated the genotypes without overlap. Its loadings recovered the speed axis without supervision: each of the five slowest motifs had a negative loading, whereas each of the four fastest motifs had a positive loading, and the loadings tracked the mean displacement of the respective motifs across the repertoire (Spearman *ρ* = 0.82, *P* = 0.007). Thus, movement speed was the dominant axis of behavioral variation in this cohort, and genotype varied along this axis. PC2 (8.4% of the variance) was unrelated to speed (*ρ* = 0.24, *P* = 0.53) and instead contrasted the three *Immobility* states (+0.63 for cluster 4 versus −0.60 and −0.34 for clusters 9 and 7, respectively). This component captured variation *within* the affected group: TG animals were 7.2 times more dispersed than WT animals along PC2. Thus, WT animals formed a tight cluster, whereas tottering animals were slower and exhibited distinct individual patterns of slowing. The low-motion states that would have been merged in a coarser vocabulary were precisely those that distinguished the TG animals from one another.

### Tottering mice hold motifs longer and sequence them more predictably

The Gantt charts suggested that TG mice used motifs differently and *remained* in them for longer, a feature that could be measured directly from the G-HMM’s window-by-window state assignments. Bout durations, defined as maximal runs of a single motif, had a markedly heavier tail in TG animals (Fig 7a). The 95th-percentile bout duration was 12.3 ± 2.0 s in TG animals versus 8.0 ± 0.0 s in WT animals (*P* = 7 × 10^−4^). The groups were completely separated, with every TG animal exceeding every WT animal (Fig 7b). No single motif accounted for this difference: it remained significant after removal of *any* one of the nine motifs (all *P <* 0.002), including the simultaneous removal of the two motifs with the longest bouts.

**Fig 7.**
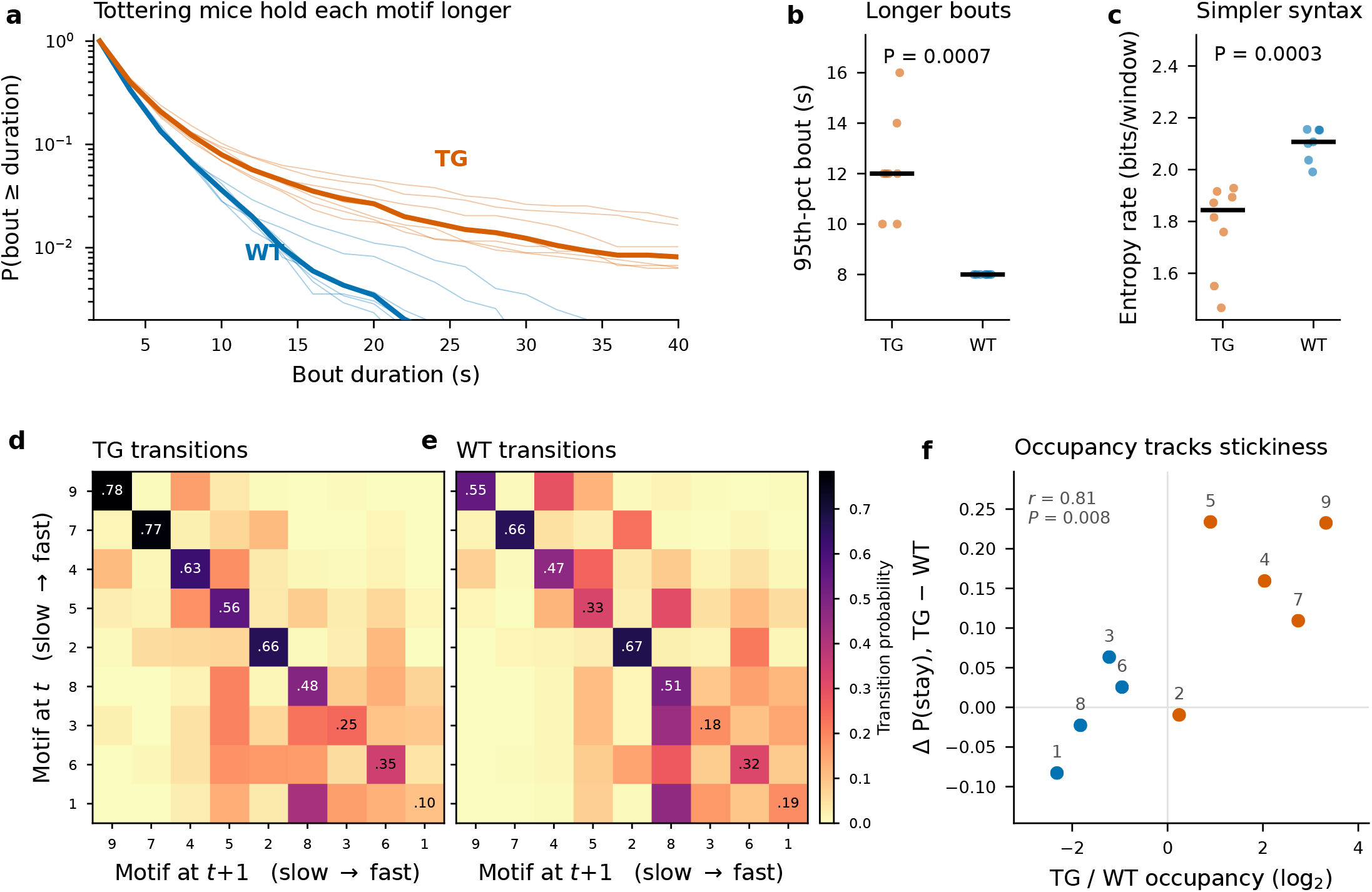
Tottering mice hold motifs longer and sequence them more predictably. All panels use the nine behavioral motifs. The artifact state (cluster 0) is excluded, and each recording is cut at the resulting gaps rather than spliced, so that no bout or transition spans an artifact. **(a)** Survival function of motif bout duration. Thin lines, individual animals; bold lines, group median (*n* = 8 TG, *n* = 7 WT). **(b)** 95th-percentile bout duration per animal (bar, median). **(c)** Entropy rate of each animal’s motif-transition matrix; lower values indicate more predictable sequencing. **(d,e)** Motif-transition matrices *P* (motif*_t_*_+1_ | motif*_t_*) pooled within genotype and displayed on a shared color scale. Motifs are ordered from slowest to fastest, as in Fig 4b, such that the low-motion states occupy the upper-left corner of each matrix; printed values are self-transition (diagonal) probabilities. **(f)** Relationship between the two axes of the phenotype, with one point per motif: the relative occupancy of the motif by TG animals (as shown in Fig 4c) is plotted against its relative persistence in TG animals. Colors indicate the predominant genotype, numbers denote cluster indices, and *r* is Pearson’s correlation across the nine motifs. *P* values in (b) and (c) were obtained using two-sided Mann–Whitney *U* tests across animals.

Because the G-HMM learns an explicit transition matrix, we could further examine whether the *sequencing* of motifs differed in addition to their duration. The entropy rate of each animal’s transition matrix, defined as the average uncertainty regarding the next motif, was lower in TG animals (1.78 ± 0.17 vs 2.10 ± 0.06 bits/window, *P* = 3 × 10^−4^; Fig 7c). The groups again showed no overlap, indicating a more restricted behavioral syntax. The transition matrices localized this difference to the diagonal (Fig 7d,e): the off-diagonal transition patterns were largely shared between genotypes (*r* = 0.83), whereas self-transitions differed 2.9 times as much. Thus, the principal difference was how long an animal remained in a motif after entering it. Persistence increased in the four slowest motifs (Δ*P* (stay) = +0.11 to +0.23) but not in the faster motifs (−0.08 to +0.06). The two axes of the phenotype therefore converged: the motifs over-occupied by TG animals were also those in which they remained longer (*r* = 0.81, *P* = 0.008; Fig 7f).

These results refine the phenotype. Tottering mice differed from their WT controls in the motifs they used, the time for which they maintained them, and the predictability with which one motif followed another. A static occupancy measure cannot capture this slowing and stereotyping of behavioral sequences. These features were accessible because the latent trajectory was modeled as a sequence rather than partitioned window by window.

### Automated behavioral metrics align with expert scores

Having shown that GESTURE uncovered a stable, data-driven phenotype, we next evaluated its biological relevance. We tested whether automatically derived behavioral metrics tracked a gold-standard, human-annotated disability score used to assess severity in this model [33]. We therefore developed two quantitative measures of behavioral divergence between the WT and TG groups based solely on the unsupervised cluster labels.

Our first metric was a ‘low-motion divergence’ measure, defined as the difference between TG and WT occupancy of the low-activity motifs (clusters 4, 5, 7, and 9). This selection was not arbitrary: these were simultaneously the four slowest states, the four most strongly over-occupied by TG animals (Fig 4c), and the four with the largest increases in self-transition probability in TG animals (Fig 7f). The metric peaked approximately 20–40 min into the session, indicating that TG mice differed most strongly from WT animals during this period (Fig 8a).

**Fig 8.**
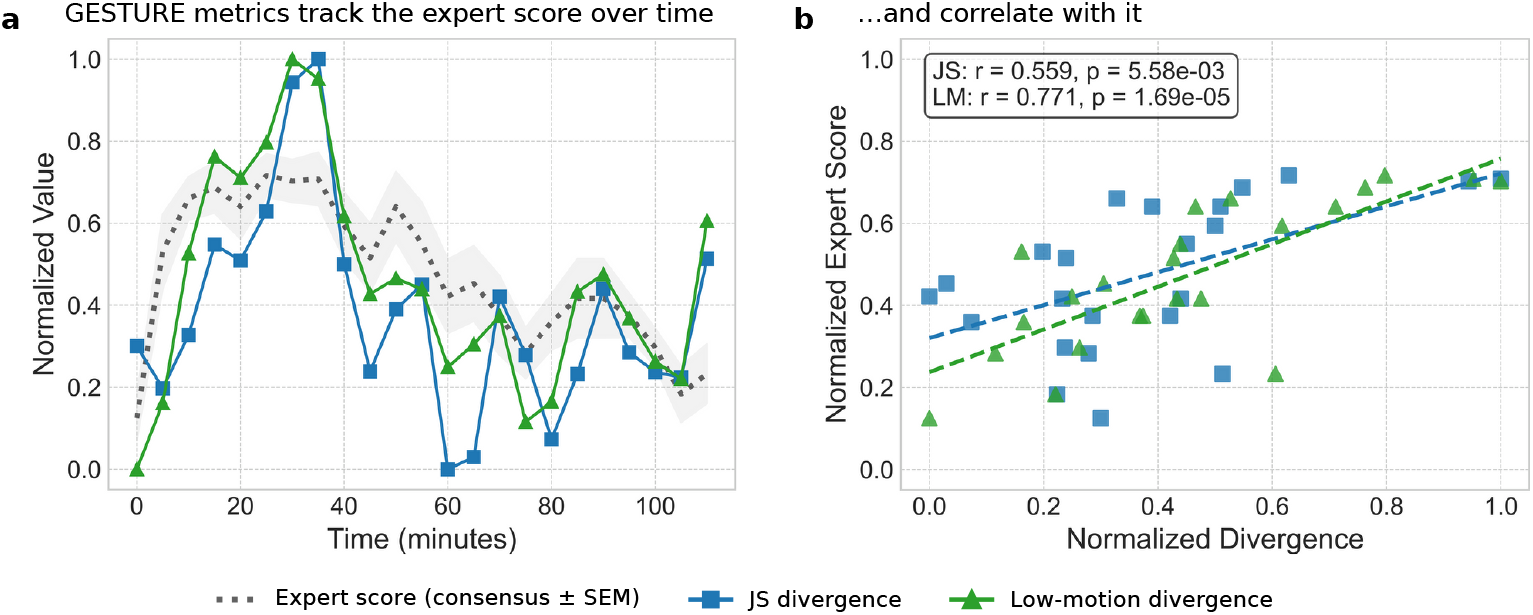
GESTURE-derived metrics track expert-annotated disability scores. **(a)** Time courses of the consensus expert-annotated disability score (dotted gray line; shaded band, SEM across animals) and the GESTURE-derived JS divergence (blue squares) and low-motion divergence (green triangles). All series are min–max normalized. **(b)** Each GESTURE-derived metric plotted against the expert consensus score, with the least-squares fit (Pearson *r*; *n* = 23 5-min bins).

The second, fully automated metric was computed as the **Jensen–Shannon (JS) divergence** between the complete 10-cluster behavioral distributions of the TG and WT groups within a sliding time window. This global measure of behavioral dissimilarity yielded a similar temporal profile and again peaked during the 20–40-min interval (Fig 8a). Pearson correlation analysis against the consensus expert scores, downsampled to 5-min intervals, revealed significant positive correlations for both low-motion divergence (*r* = 0.771, *P* = 1.69 × 10^−5^) and JS divergence (*r* = 0.559, *P* = 5.58 × 10^−3^) (Fig 8b).

Because each animal was scored independently by two trained raters, the agreement between their annotations provided a reference ceiling. The two experts agreed at *r* = 0.896 for the cohort-mean time course (ICC(2, 1) = 0.894) and at *r* = 0.870 ± 0.076 across individual animals. The animal-level correlations ranged from 0.765 to 0.974, indicating appreciable disagreement even between trained observers scoring the same animal. GESTURE’s agreement with the expert consensus (*r* = 0.771) fell within this range. These findings indicate that GESTURE-derived features provide quantitative proxies for the gold-standard, manually scored disability measure and therefore represent candidate behavioral biomarkers derived directly from raw video.

### Explainability analysis

Finally, to determine which anatomical features most strongly shaped the latent representation, we used explainability analyses to examine the encoding strategy learned by the model. When aggregated across all behavioral windows, saliency was concentrated on the trunk and limbs rather than on peripheral or cranial landmarks (Fig 9a).

**Fig 9.**
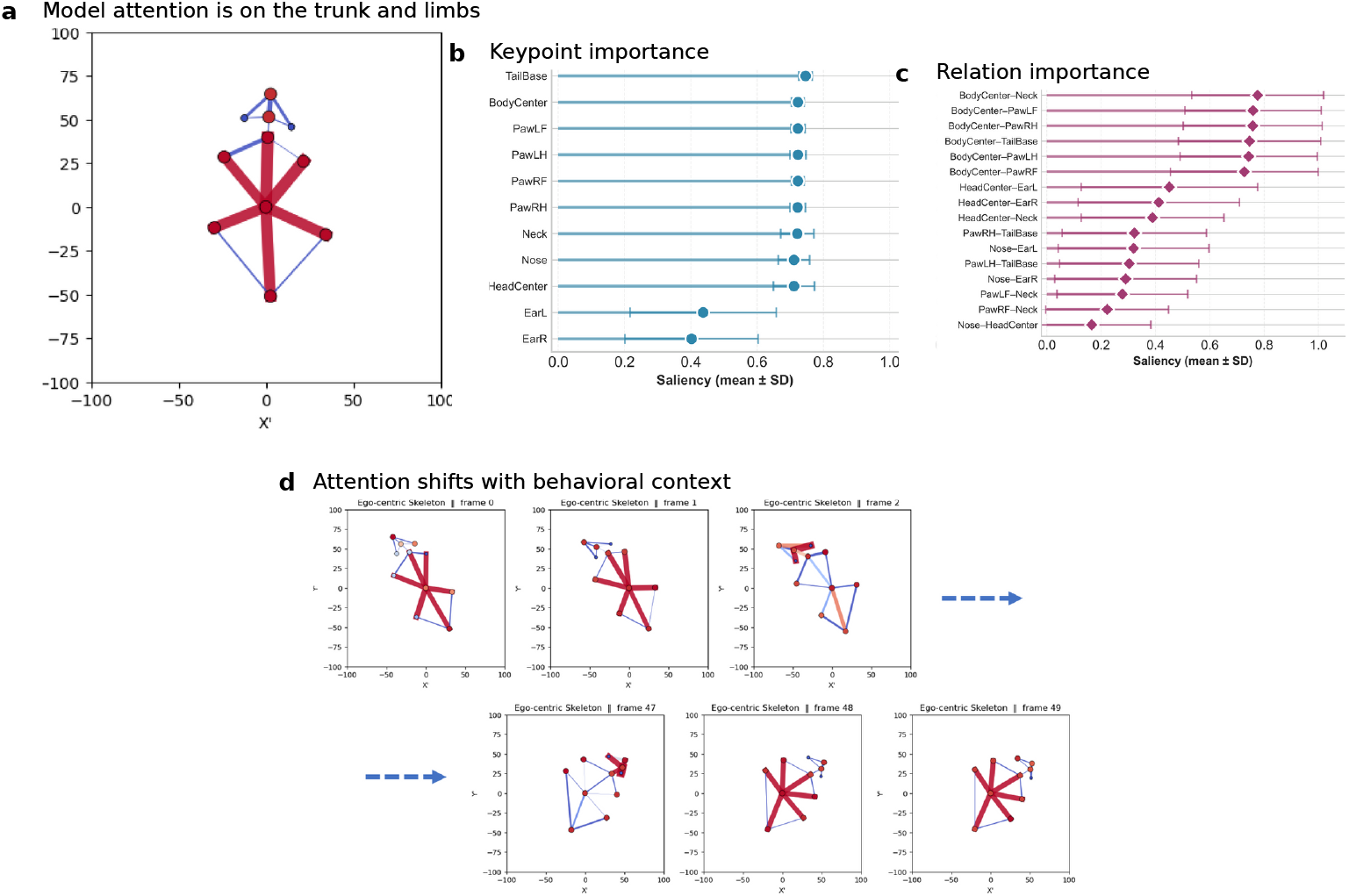
The latent representation is shaped by the animal’s core motor scaffold. **(a)** Global saliency map, aggregated over all behavioral windows. Edge importance is indicated by color and thickness; attention is concentrated on the trunk and limbs. **(b)** Node (keypoint) importance and **(c)** edge (relation) importance, shown as the mean ± SD across all windows (Eq 4) and sorted by the mean. **(d)** Frame-resolved saliency maps within a single behavioral window, showing that the model’s emphasis shifts with behavioral context: axial cues dominate at the onset of rotation (top row), whereas limb-support edges dominate during a locomotor burst (bottom row).

Quantitatively, *TailBase* had the highest mean node importance (0.746 ± 0.020), followed closely by *BodyCenter* and all four paws, which formed a low-variance second tier (*µ* ≈ 0.72, *σ <* 0.025); the ears were the least salient (Fig 9b). The most salient *relations* were similarly anchored at the trunk: *BodyCenter–Neck*, the four *BodyCenter–Paw* edges, and *BodyCenter–TailBase*. Cranial edges were less salient and substantially more variable relative to their means (Fig 9c). Thus, trunk-centered relations contributed consistently across behavioral windows, whereas head and ear saliency was context dependent.

Frame-resolved saliency maps further showed that this emphasis was dynamic rather than fixed (Fig 9d). During turning, saliency was concentrated on axial cues such as the *BodyCenter–TailBase* edge, whereas during locomotor bursts it shifted toward the limb-support edges that drive propulsion. These analyses suggest that the latent representations learned by GESTURE are shaped by biologically plausible, behavior-specific anatomical relations in the animal’s core motor scaffold rather than by idiosyncratic peripheral cues.

## Discussion

We present GESTURE, an unsupervised deep learning framework that autonomously discovers and quantifies genotype-associated behavioral structure in a mouse model of motor dysfunction. GESTURE models high-dimensional pose dynamics using an architecture that represents both spatial anatomy and temporal evolution, yielding interpretable ‘behavioral fingerprints.’ These fingerprints separated WT and TG mice without supervision (Fig 6; Fig S3 Fig). Automated divergence metrics derived from the fingerprints tracked the temporal dynamics of expert-annotated disability scores and reached agreement comparable to that between the two human raters (Fig 8). These findings support the biological relevance of the learned representation.

Modeling behavior as a *sequence* rather than as a static partition was important beyond improving label smoothness. Because the G-HMM learns an explicit transition matrix, the temporal organization of behavior becomes directly measurable. This analysis revealed a feature of the tottering phenotype that motif occupancy alone does not capture. Affected animals used low-activity motifs more often, maintained each motif substantially longer, and sequenced motifs more predictably, with a significantly lower transition entropy rate than controls (Fig 7). The phenotype therefore reflects a *slowing and stereotyping* of behavioral sequences rather than simply reduced activity. Methods that cluster behavioral windows independently of time cannot capture this distinction (Fig 2).

This work has implications for both basic neuroscience and preclinical pharmacology. High-dimensional behavioral fingerprints provide an automated, reproducible, and interpretable readout that complements traditional summary metrics and could help quantify how interventions shift an animal’s behavioral repertoire. For example, they could be used to assess whether a compound shifts a disease-associated fingerprint toward the WT distribution or changes behavioral organization in control animals.

More broadly, our results support the feasibility of deriving quantitative, interpretable behavioral signatures directly from video-based pose dynamics. Anatomically structured representation learning may therefore complement existing unsupervised behavioral pipelines by treating coordination among body parts as a direct object of analysis rather than a by-product of feature engineering or latent compression.

Two caveats constrain these conclusions. The phenotype was organized along an axis of displacement in the image plane from a single overhead view, which under-resolves rearing and fine tremor. In addition, the ethological names assigned to the motifs were based on inspection of average motion profiles and were not validated against independent annotations. The interpretability of the vocabulary is therefore supported by weaker evidence than its discriminative power.

Several directions warrant further study. First, the current implementation relies on a predefined skeletal graph. Learning graph structure from data and systematically varying graph connectivity will help distinguish architectural inductive biases from learned behavioral structure. An anatomy-defined graph is necessarily an approximation and may underrepresent task-dependent nonlocal couplings or interactions with the environment. The explainability analyses also probe the sensitivity of the latent representation to anatomical inputs but do not provide a causal account of genotype separation. Second, the present study focuses on a simplified, single-animal open-field assay. Extending the framework to 3D pose and multi-animal social interactions will be important for broader applicability. Finally, linking behavior to its neural substrates remains a key goal. The structured latent space learned by GESTURE provides a scaffold for integration with simultaneously recorded neural activity, allowing future studies to align behavioral motifs with circuit-level dynamics.

## Author contributions

Conceptualization: RQ, SF, CLE, and UR; Methodology: RQ, SF, CLE, and UR; Software: RQ, SF, and TJ; Formal analysis: RQ and SF; Investigation: RQ, EC, and KP; Data curation: RQ, SF, EC, and KP; Validation: RQ, EC, and KP; Resources: FG, CLE, and UR; Visualization: RQ; Supervision: FG, AB, CLE, and UR; Project administration: CLE and UR; Funding acquisition: UR; Writing – original draft: RQ; Writing – review and editing: all authors. All authors discussed the results and approved the final manuscript.

## Acknowledgments

We thank Nicolai Lolansen for assistance with the analysis computing environment. We thank Jesper Frank Bastlund and Benjamin Hall for valuable discussions.

## Data availability

All code required to reproduce the analyses, together with the trained model weights and the per-animal behavioral motif label sequences underlying every quantitative result reported here, is publicly available at https://github.com/HughYau/GESTURE_code.

## Supporting information

**S1 Text. Supplementary methods.**

### Pose estimation and keypoint tracking

For pose estimation, we used DeepLabCut [4], a widely adopted deep learning framework for markerless tracking of animal posture. Lundbeck researchers had trained the initial pose estimator on 300 manually labeled frames from their proprietary video archive. This pretraining provided a strong starting point, but the model had been tuned to earlier experimental setups and could not be transferred directly to the new recording environment.

The target recordings differed substantially from those obtained in standard open-field setups. The arena included a drinking pipe in the upper-right quadrant, a food area in the upper center, and a transparent rest box in the lower left, as well as altered lighting conditions and modified arena dimensions (Fig S1 Fig). These changes produced a distribution shift relative to the data on which pose estimators are typically pretrained.

**S1 Fig.**
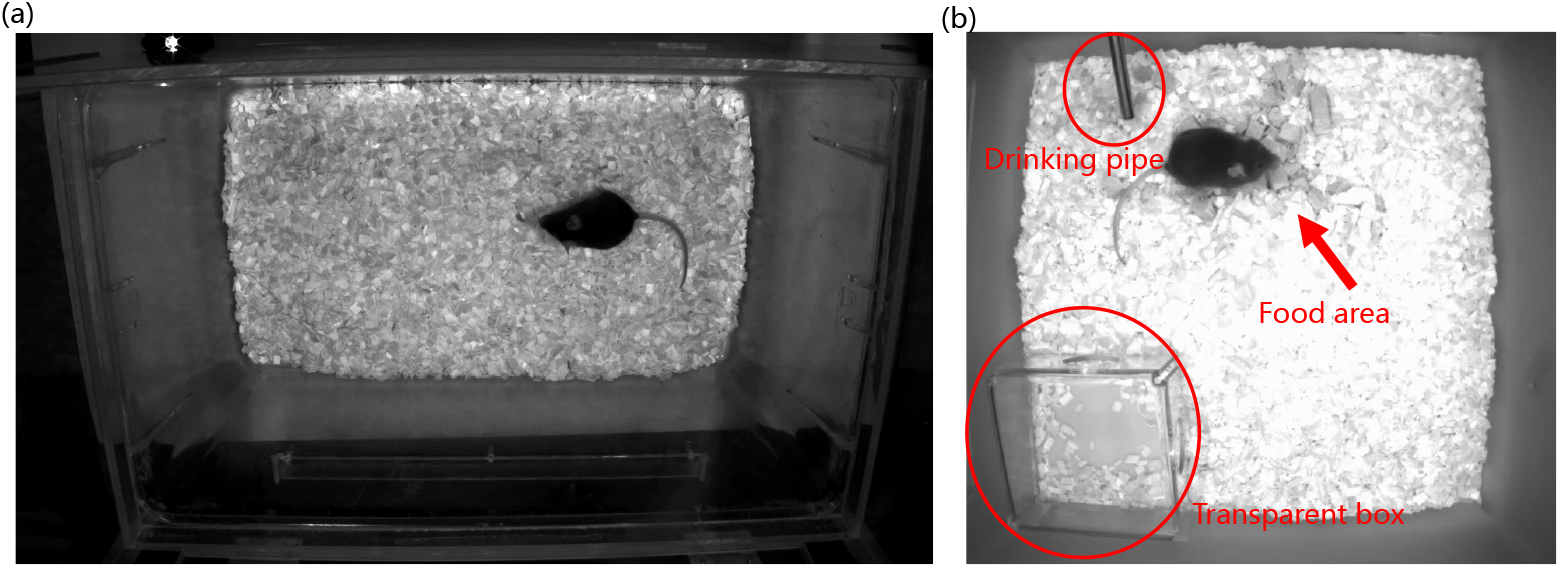
Modified arena setup in the previous (a) and target (b) datasets. Environmental structures, including a drinking pipe, food area, and transparent box, were added, and the overall scene brightness was increased.

To address this domain mismatch, we applied targeted transfer learning. We manually annotated 100 frames from the target dataset, sampling diverse mouse poses, spatial locations, and interactions with arena structures. The DeepLabCut network was then fine-tuned on this new dataset. This adaptation increased keypoint-detection likelihoods across nearly all joints and produced more spatially consistent, arena-aware predictions (Fig S2 Fig). Most keypoints achieved mean detection likelihoods above the quality-control threshold of 0.8, supporting the effectiveness of the domain-adaptation procedure (Fig S2 Figc).

**S2 Fig.**
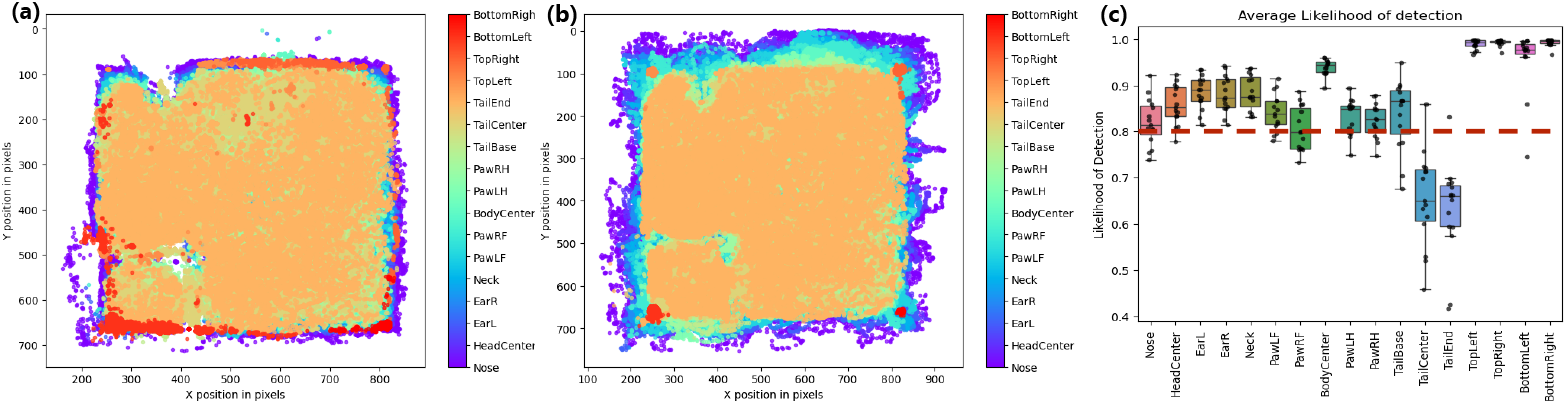
Improvement in pose estimation following domain-specific fine-tuning. Keypoint heatmaps **(a)** before and **(b)** after adaptation. **(c)** Detection-likelihood distributions for each keypoint after adaptation.

Following adaptation, each video was transformed into a temporally ordered sequence of 2D coordinates for a curated set of 11 keypoints: nose, head center, left and right ears, neck, body center, all four paws, and tail base. Each keypoint was associated with a per-frame likelihood score reflecting detection confidence. TailCenter and TailEnd, which had systematically low detection quality, were excluded from all downstream analyses because their mean likelihoods were below the 0.8 threshold. This exclusion increased the robustness of the input representation and limited the propagation of unreliable information into the model.

### Gaussian hidden Markov model

Let {*z_t_*}^*T′*^_*t*=1_ denote the sequence of latent embeddings from inlier windows, where *z_t_* ∈ R*^d^* is the VAE-encoded representation at time *t*. We model this sequence using a first-order hidden Markov model with *K* latent states. Each hidden state *s_t_* ∈ {1*, . . . , K*} emits a latent vector according to a multivariate Gaussian [29]:

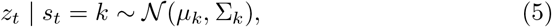

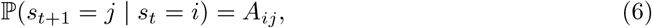

Here, *µ_k_* ∈ R*^d^* and Σ*_k_* ∈ R*^d^*^×^*^d^* denote the emission parameters of cluster *k*, and *A* ∈ R*^K^*^×^*^K^* is the state-transition matrix. This formulation jointly captures spatial similarity through Gaussian likelihoods in latent space and temporal continuity through state transitions, yielding smooth, interpretable state sequences. The model parameters Θ = {*µ_k_,* Σ*_k_, A*}^*K*^_*k*=1_ are estimated by expectation–maximization to maximize the sequence likelihood:

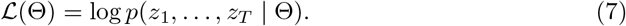

The estimated state sequence {*s_t_*}^*T′*^_*t*=1_ provides frame-level behavior annotations that preserve temporal structure, unlike static clustering methods such as *k*-means. Training the HMM on the latent sequence yields a set of recurring latent states that summarize the manifold of normal behaviors (Fig 2) and transition probabilities that model temporal progression between states. Unlike static clustering methods such as *k*-means, the HMM preserves the sequential order of windows: it groups similar latent vectors while accounting for how the corresponding clusters progress over time according to the learned dynamics.

### Determining the number of states (***K***) using BIC

We determine the number of latent states *K* from the data using the Bayesian information criterion (BIC) [34]. Specifically, we train G-HMMs across a range of candidate values and compute

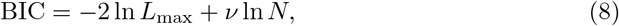

where *L*_max_ is the maximized model likelihood, *ν* is the number of parameters (which increases with *K*), and *N* is the number of data points (i.e., the total number of inlier windows when anomaly filtering is applied). BIC balances model fit and complexity by penalizing additional parameters. We select the value of *K* that yields the lowest BIC, thereby reducing the risk of choosing an unnecessarily large number of clusters that would overfit idiosyncrasies in the data.

### Statistical validation of the genotype separation

**S3 Fig.**
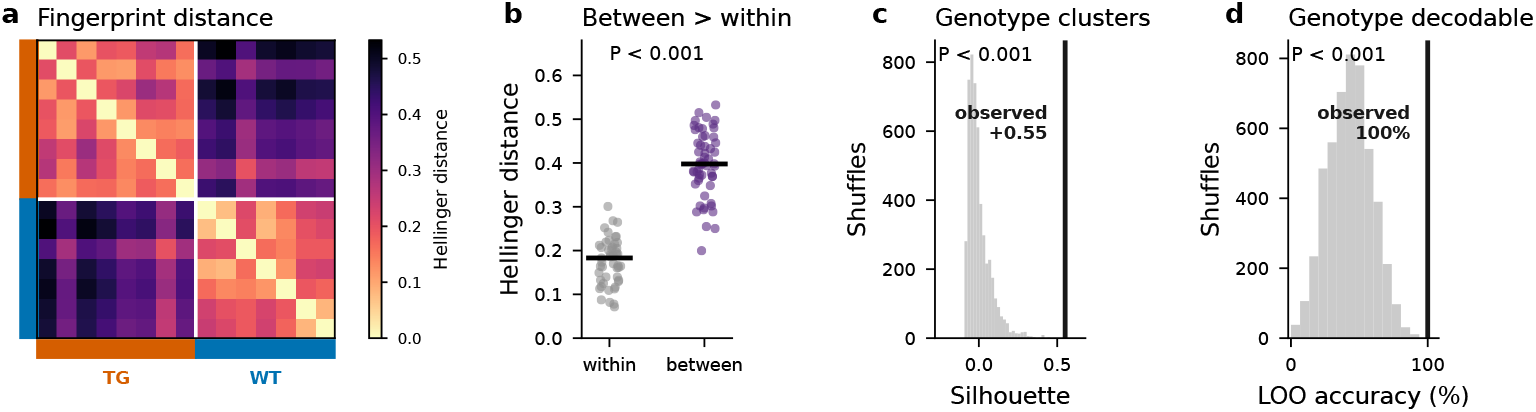
The behavioral fingerprint separates genotype. Statistical validation of the genotype separation shown in Fig 6. **(a)** Pairwise Hellinger distances between the per-animal motif-occupancy fingerprints of all 15 animals, ordered by genotype (TG, *n* = 8; WT, *n* = 7); the block structure reflects the genotype split. **(b)** Between-genotype distances exceed within-genotype distances (each point represents one animal pair; bar, median). **(c)** Silhouette of the genotype labels in fingerprint space (vertical line) compared with the label-shuffled null distribution (gray). **(d)** Leave-one-out genotype classification accuracy based on the fingerprint alone (vertical line) compared with the label-shuffled null. All *P* values were obtained using label-permutation tests. With *n* = 15 and an 8/7 split, only ^15^ = 6435 distinct labelings exist; thus, *P* ≈ 2 × 10^−4^ is the smallest attainable value.

Figure 6 depicts the separation qualitatively using a dendrogram, a distance heatmap, and a projection. Figure S3 Fig quantifies the separation directly from the per-animal motif-occupancy fingerprints and genotype labels alone.

### Zero-shot generalization to unseen animals

GESTURE is a self-supervised model trained solely on a reconstruction objective. No genotype label, behavioral annotation, or expert score enters the encoder at any point. We therefore asked whether the learned representation *transferred* to unseen animals. Six of the fifteen recordings were used to fit the encoder, whereas nine were not. These nine recordings could therefore be encoded zero-shot using a mapping that was fixed before those animals were encountered and was never exposed to any label, allowing the phenotype to be measured through the fixed representation.

The representation transferred to the unseen recordings. Among the nine unseen recordings (5 TG and 4 WT), the same leave-one-out nearest-neighbor classifier recovered genotype from the fingerprint in 9*/*9 animals. The silhouette of the genotype labels was +0.668, compared with +0.598 in the full cohort. Both sequence statistics separated the groups without overlap: the 95th-percentile bout duration was 12.8 ± 2.3 s in TG animals versus 8.0 ± 0.0 s in WT animals (*U* = 20, *P* = 0.015), and the transition entropy rate was 1.71 ± 0.20 versus 2.13 ± 0.03 bits/window (*U* = 0, *P* = 0.016). The larger *P* values relative to those for the full cohort reflect the smaller sample (*n* = 9 rather than *n* = 15), rather than a weaker effect. With nine animals split 5/4, *P* = 0.008 is the smallest value that a two-sided Mann–Whitney test can return.

## References

1. Sweis BM, Nestler EJ. Pushing the boundaries of behavioral analysis could aid psychiatric drug discovery. PLoS Biology. 2022;20(12):e3001904.

2. Vincent F, Nueda A, Lee J, Schenone M, Prunotto M, Mercola M. Phenotypic drug discovery: recent successes, lessons learned and new directions. Nature Reviews Drug Discovery. 2022;21(12):899–914.

3. Luxem K, Mocellin P, Fuhrmann F, Kürsch J, Miller SR, Palop JJ, et al. Identifying behavioral structure from deep variational embeddings of animal motion. Communications Biology. 2022;5:1267. Available from: 10.1038/s42003-022-04080-7. doi:10.1038/s42003-022-04080-7.

4. Mathis A, Mamidanna P, Cury KM, Abe T, Murthy VN, Mathis MW, et al. DeepLabCut: markerless pose estimation of user-defined body parts with deep learning. Nature Neuroscience. 2018. Available from: https://www.nature.com/articles/s41593-018-0209-y.

5. Tara E, Vitenzon A, Hess E, Khodakhah K. Aberrant cerebellar Purkinje cell activity as the cause of motor attacks in a mouse model of episodic ataxia type 2. Disease models & mechanisms. 2018;11(9):dmm034181.

6. Ebner TJ, Carter RE, Chen G. Tottering mouse. In: Handbook of the cerebellum and cerebellar disorders. Springer; 2021. p. 1709–32.

7. Anderson DJ, Perona P. Toward a Science of Computational Ethology. Neuron. 2014;84(1):18–31. Available from: 10.1016/j.neuron.2014.09.005. doi:10.1016/j.neuron.2014.09.005.

8. Datta SR, Anderson DJ, Branson K, Perona P, Leifer A. Computational Neuroethology: A Call to Action. Neuron. 2019;104(1):11–24. Available from: 10.1016/j.neuron.2019.09.038. doi:10.1016/j.neuron.2019.09.038.

9. Perez M, Toler-Franklin C. CNN-Based Action Recognition and Pose Estimation for Classifying Animal Behavior from Videos: A Survey. arXiv preprint arXiv:230106187. 2023. Available from: http://arxiv.org/abs/2301.06187. arXiv:2301.06187. doi:10.48550/arXiv.2301.06187.

10. Berman GJ, Choi DM, Bialek W, Shaevitz JW. Mapping the Stereotyped Behaviour of Freely Moving Fruit Flies. Journal of The Royal Society Interface. 2014;11(99):20140672. Available from: https://royalsocietypublishing.org/doi/10.1098/rsif.2014.0672. doi:10.1098/rsif.2014.0672.

11. Wiltschko AB, Tsukahara T, Zeine A, Anyoha R, Gillis WF, Markowitz JE, et al. Revealing the Structure of Pharmacobehavioral Space through Motion Sequencing. Nature Neuroscience. 2020 Nov;23(11):1433–43. doi:10.1038/s41593-020-00706-3.

12. Lin S, Gillis WF, Weinreb C, Zeine A, Jones SC, Robinson EM, et al. Characterizing the Structure of Mouse Behavior Using Motion Sequencing. Nature Protocols. 2024 Nov;19(11):3242–91. doi:10.1038/s41596-024-01015-w.

13. Stephens GJ, Johnson-Kerner B, Bialek W, Ryu WS. Dimensionality and dynamics in the behavior of C. elegans. PLoS computational biology. 2008;4(4):e1000028.

14. Nath T, Mathis A, Chen AC, Patel A, Bethge M, Mathis MW. Using DeepLabCut for 3D markerless pose estimation across species and behaviors. Nature Protocols. 2019. Available from: 10.1038/s41596-019-0176-0.

15. Pereira TD, Tabris N, Matsliah A, Turner DM, Li J, Ravindranath S, et al. SLEAP: A deep learning system for multi-animal pose tracking. Nature methods. 2022;19(4):486–95.

16. Hsu AI, Yttri EA. B-SOiD, an Open-Source Unsupervised Algorithm for Identification and Fast Prediction of Behaviors. Nature Communications. 2021;12(1):5188. Available from: https://www.nature.com/articles/s41467-021-25420-x. doi:10.1038/s41467-021-25420-x.

17. Batty E, Whiteway M, Saxena S, Biderman D, Abe T, Musall S, et al. BehaveNet: nonlinear embedding and Bayesian neural decoding of behavioral videos. Advances in neural information processing systems. 2019;32.

18. Weinreb C, Pearl JE, Lin S, Osman MAM, Zhang L, Annapragada S, et al. Keypoint-MoSeq: Parsing Behavior by Linking Point Tracking to Pose Dynamics. Nature Methods. 2024 Jul;21(7):1329–39. doi:10.1038/s41592-024-02318-2.

19. Yan S, Xiong Y, Lin D. Spatial Temporal Graph Convolutional Networks for Skeleton-Based Action Recognition. In: Proceedings of the AAAI Conference on Artificial Intelligence. vol. 32; 2018. Available from: 10.1609/aaai.v32i1.12328. doi:10.1609/aaai.v32i1.12328.

20. Wang Y, Sun Y, Liu Z, Sarma SE, Bronstein MM, Solomon JM. Dynamic graph cnn for learning on point clouds. ACM Transactions on Graphics (tog). 2019;38(5):1–12. Available from: 10.1145/3326362. doi:10.1145/3326362.

21. Bordes J, Miranda L, Reinhardt M, Narayan S, Hartmann J, Newman EL, et al. Automatically Annotated Motion Tracking Identifies a Distinct Social Behavioral Profile Following Chronic Social Defeat Stress. Nature Communications. 2023;14(1):4319. Available from: https://www.nature.com/articles/s41467-023-40040-3. doi:10.1038/s41467-023-40040-3.

22. Mullen PN, Bowlby B, Armstrong HC, Gray A, Zwart MF. PoseR: a deep learning toolbox for classifying animal behaviour. Open Biology. 2026;16(1). Available from: 10.1098/rsob.250322. doi:10.1098/rsob.250322.

23. Fletcher CF, Lutz CM, O’Sullivan TN, Shaughnessy JD, Hawkes R, Frankel WN, et al. Absence epilepsy in tottering mutant mice is associated with calcium channel defects. Cell. 1996;87(4):607–17. Available from: 10.1016/S0092-8674(00)81381-1. doi:10.1016/S0092-8674(00)81381-1.

24. Savitzky A, Golay MJ. Smoothing and differentiation of data by simplified least squares procedures. Analytical chemistry. 1964;36(8):1627–39.

25. Cho K, van Merriënboer B, Gulcehre C, Bahdanau D, Bougares F, Schwenk H, et al. Learning Phrase Representations using RNN Encoder–Decoder for Statistical Machine Translation. In: Proceedings of the 2014 Conference on Empirical Methods in Natural Language Processing (EMNLP); 2014. p. 1724–34. doi:10.3115/v1/D14-1179.

26. Schuster M, Paliwal KK. Bidirectional recurrent neural networks. IEEE Transactions on Signal Processing. 1997;45(11):2673–81. doi:10.1109/78.650093.

27. Higgins I, Matthey L, Pal A, Burgess C, Glorot X, Botvinick M, et al. beta-VAE: Learning basic visual concepts with a constrained variational framework. In: International conference on learning representations; 2017. .

28. Burgess CP, Higgins I, Pal A, Matthey L, Watters N, Desjardins G, et al. Understanding disentangling in *β*-VAE. arXiv preprint arXiv:180403599. 2018.

29. Rabiner LR. A tutorial on hidden Markov models and selected applications in speech recognition. Proceedings of the IEEE. 1989;77(2):257–86.

30. Ying Z, Bourgeois D, You J, Zitnik M, Leskovec J. Gnnexplainer: Generating explanations for graph neural networks. Advances in neural information processing systems. 2019;32.

31. Ebbesen CL, Froemke RC. Automatic mapping of multiplexed social receptive fields by deep learning and GPU-accelerated 3D videography. Nature Communications. 2022;13(1):593. Available from: https://www.nature.com/articles/s41467-022-28153-7. doi:10.1038/s41467-022-28153-7.

32. McInnes L, Healy J, Melville J. Umap: Uniform manifold approximation and projection for dimension reduction. arXiv preprint arXiv:180203426. 2018.

33. Jinnah HA, Sepkuty JP, Ho T, Yitta S, Drew T, Rothstein JD, et al. Calcium channel agonists and dystonia in the mouse. Movement Disorders. 2000;15(3):542–51. doi:10.1002/1531-8257(200005)15:3¡542::AID-MDS1019¿3.0.CO;2-2.

34. Schwarz GE. Estimating the dimension of a model. The Annals of Statistics. 1978;6(2):461–4.

